# Marine Ecosystems as Complex Adaptive Systems: Emergent Patterns, Critical Transitions, and Public Goods

**DOI:** 10.1101/056838

**Authors:** George I. Hagstrom, Simon A. Levin

## Abstract

Complex adaptive systems provides a unified framework for explaining ecosystem phenomena. In the past twenty years, complex adaptive systems has been sharpened from an abstract concept into a series of tools that can be used to solve concrete problems. These advances have been led by the development of new techniques for coupling ecological and evolutionary dynamics, for integrating dynamics across multiple scales of organization, and for using data to infer the complex interactions among different components of ecological systems. Focusing on the development and usage of these new methods, we explore how they have led to an improved understanding of three universal features of complex adaptive systems, emergent patterns; tipping points and critical phenomena; and cooperative behavior. We restrict our attention primarily to marine ecosystems, which provide numerous successful examples of the application of complex adaptive systems. Many of these are currently undergoing dramatic changes due to anthropogenic perturbations, and we take the opportunity to discuss how complex adaptive systems can be used to improve the management of public goods and to better preserve critical ecosystem services.

## 1 Introduction

Twenty years ago, as Ecosystems was setting out on its path to become one of the leading-edge outlets for progress in ecosystems research, it featured a special issue on viewing ecosystems as complex adaptive systems (Hartvigsen et al., 1998, Levin, 1998). Since that time, this perspective has seen impressive development, from both theoretical and applied perspectives (Filotas et al., 2014, Messier et al., 2015, Scheffer et al., 2012). In this paper, we will focus on three features of complex adaptive systems, through the lens of marine ecosystems. Complex adaptive systems are characterized first of all by emergent patterns, like food-web structure and nutrient cycling, that characterize community and ecosystem patterns at the macroscopic levels, but that to a large extent are byproducts of evolution at lower levels of organization. Alternative stable outcomes are hence possible (Carpenter et al., 2001, Jacob, 1977, Staver et al., 2011a,b), as are system flips from one basin of attraction to another or even the loss of a basin of attraction as the result of slow variable changes (Carpenter et al., 1999, Scheffer, 2009, Steele, 1998). Two fundamental and interrelated challenges are therefore to understand the relationships among phenomena at diverse scales of time, space and organizational complexity, developing scaling laws and moment-closure techniques; and to resolve the public-goods and common-pool-resource conflicts that are unavoidable. These challenges arise in the description and management of any complex-adaptive ecosystem, but for definiteness we explore them primarily in marine ecosystems.

The emergence of pattern has been a theme in theoretical science across all fields for many decades; indeed, Darwin, in the Origin of Species (Darwin, 1858) and other work, was endeavoring to understand the emergence of pattern in the tangled bank of interacting species, and indeed in the very emergence of species as operational taxonomic units. Spatial pattern has been of particular interest in fields from physics and chemistry to developmental biology to ecosystems, where regular patterning emerges from local interactions. Indeed, although in some cases the pattern that results is an epiphenomenon, once it arises it is fair game for evolution that feeds back to influence the processes that gave rise to it. In developmental biology, the feedback can be encoded genetically, producing heritable changes that result in the improvement of the end product. But even in physical systems, like water flow across landscapes, local processes may be modified through feedbacks to alter macroscopic outcomes (Rodriguez-Iturbe and Rinaldo, 1997)

In some cases, emergence is a constructive process, producing persistent structures through processes of self-organization, with the outcomes potentially uncertain and influenced by historical events, including initial conditions (Arthur, 1994, Jacob, 1977). Systems hence may exhibit alternative stable states and hysteresis, or even more complicated asymptotic behaviors. Where this occurs, attention falls upon the robustness and resilience of particular configurations, and the potential for sudden shifts from one to the other (Holling, 1973, Levin et al., 1998, Levin and Lubchenco, 2008, Steele, 1998), as well as upon early warning indicators of impending system flips. (Lenton, 2011, Sche er, 2009, Scheffer et al., 2012). Particularly evident in many such cases is that dynamics play out on multiple scales of space, time and organizational complexity, and that the interplay among scales presents fundamental modeling challenges (Levin, 1992).

Emergence and phase transitions are well-known and well-described in physical systems, and techniques like renormalization groups and statistical mechanics more generally have provided powerful ways to relate microscopic phenomena to macroscopic outcomes. Simple percolation and other such models elucidate the emergent dynamics inherent in local rules. In fluid mechanics, moment closure techniques allow one to go from individual-based Lagrangian descriptions to continuum Eulerian ones, and back, and extensions have been applied with success to collective motion in animal populations (Couzin et al., 2005, Flierl et al., 1999, Grünbaum and Okubo, 1994, Viswanathan et al., 2011). The emergence of stable spatial structures, from animal coat patterns to landscape vegetation, are well-described in the language of reaction-diffusion equations (Murray, 2002), and advances in computation have made agent-based models a powerful tool (Couzin et al., 2005, Pacala and Silander, 1985).

We view the interplay between processes on different scales as the central issue in dealing with complex adaptive systems: the emergence of regular pattern; the potential for system flips, and the inevitable conflicts that arise between the forces that drive individual behavior and the consequences for the systems of which they are parts. These create challenges for description, as well as for management (Arrow et al., 2014, Levin et al., 2013). Layered above all of these is the need to scale from the individual to the system, from the local to the global, and from the short-term to the long-term. Hence, these issues form the foci of this paper, the necessary next steps in moving from the abstract perspectives of 20 years ago to dealing with real systems.

## 2 Emergent Patterns in Ocean Ecosystems

The physicist Philip Anderson wrote that “more is different” (Anderson, 1972), highlighting the essential importance of emergent paterns for complex adaptive systems. Emergent patterns are large-scale structures or regularities that arise due to interactions that take place on small scales (Levin, 1998), and feed back to influence those small-scale interactions. In ecological systems, emergent patterns involve the coupling and coevolution of organisms and their environment: large-scale patterns of ecological properties such as trophic structure, nutrient usage, or collective motion are generated by small-scale interactions, but also influence the further development of those small-scale interactions and cause slow changes to the physical and chemical environment as well as to the organisms themselves. These processes have shaped the biosphere of the Earth, oxygenating the atmosphere and oceans, altering the temperature and climate, and creating new niches for life to occupy (Falkowski et al., 2008).

Complex adaptive systems provided a qualitative framework for understanding emergent patterns in ecology and evolution, but initially this framework could not be used to make quantitative predictions, preventing its application to concrete problems. In order to use the theory of complex adaptive systems fully, ecologists adapted methods from statistical physics and fluid dynamics, enabling them to calculate phenomena across scales, and invented techniques to couple ecological and evolutionary dynamics. These methods have been successfully applied to many problems, elevating complex adaptive systems from an abstract concept to a tool for deriving quantitative insight into ecological systems. Here we illustrate the use of these techniques to understand emergent patterns in collective behavior and marine biogeography.

One of the most remarkable phenomena in ecology is the ability of animal groups to engage in complex, coordinated behaviors mediated only through local interactions among group-members (Couzin and Krause, 2003). Group behaviors are emergent phenomena, arising from small-scale interactions and influencing both the fitness of individuals and the states of their ecosystem and environment. To understand how collective behaviors emerge, ecologists took inspiration from statistical physicists, who solved the similar problem of understanding the bulk properties of matter from the interactions of neighbouring atoms and molecules (Chapman and Cowling, 1970). In particular, techniques from kinetic theory, fluid mechanics, and phase transitions have been successfully applied to understand the collective behavior of animal groups (Czirók and Vicsek, 2001, Flierl et al., 1999, Grünbaum, 1998). The starting point for all of these methods is a description of the interactions among individuals. *An individual-level* or *Lagrangian* description of an animal group represents how each animal changes its state as a function of the state of the environment and of the other individuals, which can be formulated mathematically as stochastic differential equations:

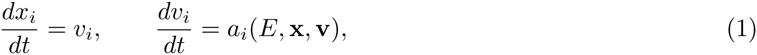

where x and v are vectors whose components *x*_*i*_ and *v*_*i*_ represent the position and velocity of the *i*th individual in an animal group, *E* represents the state of the environment, and the function *a*_*i*_ encodes the behavior of the *i*th individual and generally has a stochastic component. The individual-level variables give a precise description of the group, but are unsuitable for expressing phenomena involving large length scales or large numbers of individuals. Collective behaviors become apparent in descriptions based on *Eulerian* variables, which are the statistical moments of the probability distribution *f*(*x*, *v*) over individual position and velocity:

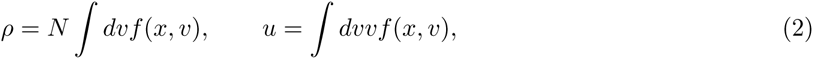

here *ρ* is the spatial density of individuals and *u* is the mean velocity. To derive equations for the Eulerian variables we take moments of equation 1, which introduces a closure problem: the evolution equation for each moment depends on the value of higher-statistical moments, leading to a hierarchy of interdependent equations. This hierarchy must be truncated and closed by approximating higher-order moments in terms of lower-order ones. These approximations are derived using assumptions about the underlying biological system (Flierl et al., 1999, Norberg et al., 2001). The parameters describing individual-level properties appear in non-trivial combinations in the resulting equations, allowing complex, large-scale dynamics to emerge from the details of the microscopic interactions. The most dramatic example of this is a phase transition, in which large-scale properties undergo sudden, sometimes discontinuous changes under small variations of the individual-level parameters.

Early efforts to understand the emergence of collective behavior modelled animal groups as following behavioral rules similar to interaction laws in physics. The sacrifice of biological realism for analytical simplicity was successful because it predicted emergent behaviors that were observed in biological ensembles of real organisms. Two separate sets of studies of the collective motion of abstract agents showed that a transition from undirected motion to collective motion could arise through an interplay of alignment, navigational noise and velocity (Toner and Tu, 1995, Vicsek et al., 1995). As alignment forces increase, noise decreases, or speed decreases, the velocities of the agents undergo a phase transition from a disordered state to an aligned state. Subsequently, lab and field measurements of different species of schooling fish identified various distinct behavioral states such as swarming, milling, and polarized swimming (Katz et al., 2011), and theoretical methods were used to explain the states in terms of individual-level behaviors (Tunstrøm et al., 2013). A parallel series of work initiated by Flierl, Grunbaum, and others (Flierl et al., 1999, Grünbaum, 1994) used moment closures to explore the factors governing the mobility of groups, the interaction between individual behavior and fluid flow, and group size distributions among different types of motile marine organisms. These papers derived advection-diffusion equations and more complicated continuum equations containing integral terms, to model social organisms and their interaction with with patchy resource distributions (Grünbaum, 1998) and which also could be compared to real data. Development of more advanced methods for tracking animals has led to increased adoption of the moment-closure modelling framework, both in marine and terrestrial ecology, and has catalyzed the recent synthesis in movement ecology (Mueller and Fagan, 2008).

Computer vision enables precision tracking of the behavior of individuals within groups, allowing for the generation of time-series of individual positions and responses to both environmental conditions and social cues (Berdahl et al., 2013, Cavagna et al., 2010). These developments have led to models with biologicaly realistic interaction laws, incorporating social interaction networks inferred from data and opening up the possibility of analyzing the emergent properties of social networks for many types of social animals, including schooling fish (Rosenthal et al., 2015). These observations have shown that under some conditions behavioral rules are better described by topological interactions, in which each individual pays attention to a fixed number of nearest neighbors, than by metric interactions in which an individual pays attention to all neighbours within a fixed distance (Berdahl et al., 2013, Brush et al., 2016, Cavagna et al., 2010). Observed interaction networks have a structure that maximizes information transfer as well as network robustness (Young et al., 2013), placing animal groups at the edge of phase transitions (Bialek et al., 2012, Hein et al., 2015). The evolution of collective behavior has also been investigated using evolutionary simulations, in which a large number of behaviorally heterogeneous agents searched a noisy environment for scarce patches of resources using a combination of social and environmental information. Evolved populations gained the ability to collectively detect gradients and to track stochastically moving resource patches, demonstrating a mechanism by which collective behaviors can evolve via natural selection on selfish individuals (Hein et al., 2015). The shift to studies of biologically realistic models has also led to the modification of the moment closure techniques and the theory of non-equilibrium phase transitions, allowing for the extension of theoretical methods to these new cases. The speed of technological advancement in movement ecology is accelerating, and complex adaptive systems will continue to provide the tools that create ecological insight from the new data that will soon be available.

Earth’s environment has been shaped by the interplay between ecological and evolutionary processes, including coevolutionary feedbacks between organisms and their environments (Falkowski, 1997, Falkowski et al., 2008). Understanding how adaptive processes influence emergent patterns is one of the grand challenges of modern science, with applications not only to ecology and biogeochemistry, but also to human society and behavior. It has been proposed that to solve these problems we need a “statistical mechanics of types” (Levin, 1992) to model the appearance of new phenotypes and behavioral strategies. This proposal motivated the development of theoretical tools to model the evolutionary dynamics underlying complex adaptive systems, combining ideas from a multitude of fields, including game theory, population genetics, statistical physics, and physiology. Efforts to apply these new ideas to problems involving adaptation and coevolution have intensified in recent years, and are key steps towards fulfillment of the great promise of complex adaptive systems.

Eco-evolutionary dynamics involve two important ingredients: a representation of the space of possible phenotypes and a model of the ecological competition among those phenotypes. Trait-based models (Litch-man and Klausmeier, 2008, Shuter, 1979) represent organisms as distinct phenotypes instead of species, parametrizing them as points in a physiological trait-space, providing a means of determining the fitness of each phenotype as a function of the environment. A realistic trait space requires a quantitative formulation of the trade-offs among different traits, which are determined from physical and chemical constraints and physiological measurements. For instance, organism size is a fundamental trait that directly influences numerous physiological processes, including metabolic rate, swimming speed, nutrient uptake, and trophic interactions (Andersen et al., 2016). By providing a means of calculating the fitness of an organism with a given set of traits, a trait-based model provides a set of equations governing the population dynamics in trait space. To incorporate evolution, mechanisms for introducing new types, such as random mutation or immigration, must be included in the population dynamics, introducing an evolutionary time-scale. A modelling framework called adaptive dynamics (Diekmann et al., 2004, Geritz et al., 1997) applies when the ecological time-scale is faster than the evolutionary time-scale, leading to a scenario in which there is a dominant type that is repeatedly invaded by nearby mutants. Adaptive dynamics defines an object called the selection function, written *s*_*x*_(*y*), as the long term growth rate of an organism with trait y in a population dominated by trait x. The gradient of *s*_*x*_(*y*) determines the direction of evolution in trait space, and critical points of the selection function correspond to special types, such as *evolutionarily stable strategies*, which are locally uninvasible; *convergence stable strategies*, which are local attractors of the evolutionary dynamics; and *branching points*, which lead to speciation. Adaptive dynamics is particularly powerful because it can give exact analytical results demonstrating the circumstances under which many important eco-evolutionary phenomena can occur.

Adaptive dynamics has been used to explore eco-evolutionary dynamics and emergent patterns in a wide variety of ecosystems, resulting in many surprising predictions that would have been difficult to produce with traditional, non-evolutionary models. Sheffer and colleagues (Sheffer et al., 2015) used adaptive dynamics to explain the global distribution of facultative and obligate nitrogen-fixing trees, and in particular the long-standing mystery about why nitrogen-fixing trees dominate nitrogen replete tropical forests but are rare in nitrogen-poor temperate forests. Using a one-dimensional trait space that represented an investment into nitrogen fixation, the adaptive dynamics model predicted the coexistence of facultative nitrogen fixers with non-fixers in forests with high rates of disturbance and transient periods of nitrogen de cit. Such coexistence was impossible in low-disturbance, temperate forests, in which there was instead a successional cycle involving a transition from obligate nitrogen fixers to non-fixers. Despite having a small number of parameters, this model was able to predict global emergent patterns and also gave accurate quantitative estimates of terrestrial rates of nitrogen fixation. Adaptive dynamics has been applied to a variety of different problems, and has also yielded insights into fisheries management (Heino, 1998) and phytoplankton functional response (Lomas et al., 2014).

When ecological and evolutionary time-scales are the same it is necessary to consider the full population distribution in trait space. Trait-based models formulate the full population dynamics with an equation for the net growth rate at each point in trait space:

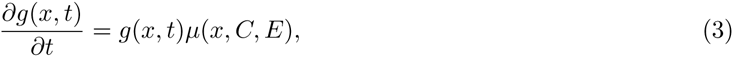

where *g*(*x*, *t*) is the population density in trait space, *x* is the trait value, *μ* is the net growth (or death) rate, *E* describes the environment, and *C* is the total biomass. Equation 3 is usually too complex to be solved directly, and there are numerous ways in which it can be approximated or simplified. The moment closure methods introduced in equation 2 can be applied to equation 3, which leads to reduced equations for the mean and covariances of the trait distribution g. A standard closure for these equations in a one-dimensional trait space is (Norberg et al., 2001):

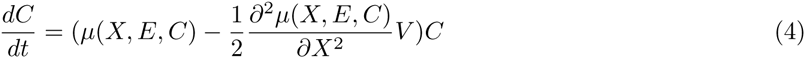

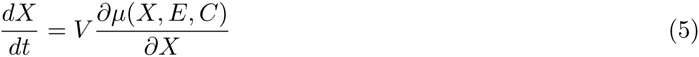

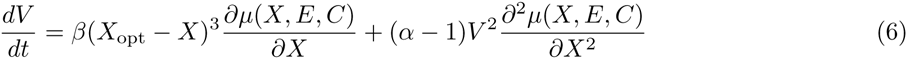

Here 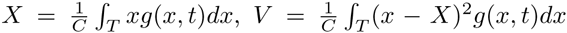, the numbers *α* and *β* are coefficients that are calculated from the trait-based model during the moment closure procedure, and *X*_opt_ is the trait-value that maximizes the fitness *μ* for the environment *E*. Equation 4 implies that the biomass increases at a rate proportional to the mean growth rate, but with a lag proportional to the trait variance and the convexity of the fitness function *μ*, so that diversity can negatively impact population growth. Equation 5 implies that the mean trait value changes in the direction of the gradient of *μ*, which pushes the trait towards *X*_opt_, the trait which maximizes *μ* in the environment *E*. The moment equations predict that the mean trait will approach the value that optimizes the fitness *μ*, and the trait dynamics can be drastically simplified by assuming *X* = *X*_opt_, implying rapid adaptation of the population to the current environment. Approaches based on moment equations or optimality principles are poor if there is no unique optimum of the fitness function, if multiple phenotypes are expected to coexist, or if the trait space is very high-dimensional and it is unclear how to approximate the trait-dynamics. In these cases, emergent ecosystems can be simulated by seeding the population model with phenotypes selected at random from the trait space.

All of the different methods for solving equation 3 have been used to elucidate the environmental determinants of phytoplankton biogeography. Follows et. al. (Follows et al., 2007) used random-sampling from a trait space parametrized by laboratory measurements of phytoplankton functional responses, defining phytoplankton by their light and temperature preferences, their nutrient half-saturation constants, and their maximum growth rates. The relative values of these traits were constrained strongly by data, which predicts a trade-off between maximum growth rate and competitiveness for nutrients (roughly equivalent to the trade-offs between r-strategists and K-strategists). 70 phenotypes were chosen to seed a general circulation model, and model community biogeography was compared to observed phytoplankton biogeography. The similarities in community structure were remarkable, verifying that phytoplankton biogeography can emerge from competition along a continuum of r-strategists and K-strategists (Armstrong, 1994, Kiørboe, 1993, Margalef et al., 1979) and predicting that the absence of *Prochlorococcus*, a picophytoplankton that thrives in ultra-oligotrophic gyres, from sub 15 degree waters is due to high levels of nutrient loading in cold water rather than an inability for *Prochloroccocus* to physiologically adapt to low temperatures. This landmark study has inspired a host of additional work on global trait biogeography, include predictions of the global distribution of mixotrophy (Ward and Follows, 2016) and food-web size structure (Clark et al., 2013, Ward et al., 2012).

Phytoplankton elemental stoichiometry is a particularly interesting trait from the point of view of complex adaptive systems because it couples the cycling of carbon and inorganic nutrients in the ocean, mediating co-evolutionary feedbacks between resource supply, phytoplankton resource ratios, and primary productivity (Galbraith and Martiny, 2015, Hagstrom et al., 2016, Tyrrell, 1999). The interaction of phytoplankton elemental stoichiometry with marine biogeochemistry and primary productivity is strongly influenced by nitrogen fixation, which couples the deep ocean nitrate inventory to the rate of phosphorus or iron supply to the ocean (Hagstrom et al., 2016, Tyrrell, 1999). These studies suggest that understanding the environmental drivers of phytoplankton stoichiometry is crucial for having a proper understanding of primary productivity and biogeochemical cycling in the ocean. Trait-based models of phytoplankton stoichiometry have been instrumental in revealing the environmental drivers of phytoplankton resource ratios. Using a model that linked the investments of phytoplankton into resource gathering and protein synthesis to both phytoplankton fitness and phytoplankton N:P ratios, Klausmeier et. al. (Klausmeier et al., 2004) showed that the optimal phytoplankton strategy in an oligotrophic environment is to have large investments in nitrogen-rich proteins and chloroplasts and thus a high N:P ratio, whereas the optimal phytoplankton strategy in a eutrophic environment is to have a large number of ribosomes and thus a low N:P ratio. This suggested that the Red eld ratio (the mean N:P ratio of phytoplankton in the ocean) is not a global physiological optimum, as was previously believed (Redfield, 1958), but instead emerges from a combination of high N:P phytoplankton in oligotrophic gyres and low N:P phytoplankton in upwelling regions. This prediction was confirmed by observational studies of the stoichiometry of particulate organic matter (Martiny et al., 2013) and of inorganic nutrient tracers (Teng et al., 2014). Since then, more precise trait-based models of stoichiometry have been developed, which used both the optimality approach and a refined random-sampling method (Clark et al., 2013), allowing for the prediction of the separate influences of light levels, nutrient supply, and temperature on phytoplankton N:P ratios (Daines et al., 2014, Toseland et al., 2013). These predictions have led to theoretical models suggesting that both global warming (Toseland et al., 2013) and increased nutrient depletion in oligotrophic gyres (Galbraith and Martiny, 2015) could led to large drawdowns in atmospheric CO2. The fact that N:P ratios show systematic variation in the ocean implies that there can be complex feedbacks between marine phytoplankton and biogeochemical cycles. Primary productivity, nutrient limitation patterns, and the rate of nitrogen fixation depend on the relative rates of nutrient supply but also on the nutrient stoichiometry. In order to fully understand the co-evolution of stoichiometry and environment, trait-based stoichiometric models need to be extended to better predict the iron requirements of phytoplankton and to predict the level of luxury storage of non-limiting nutrients.

Challenges remain to make the best use of the insights derived from eco-evolutionary models of ecosystems. Random-sampling approaches to trait-based models are widely known, but require high levels of computational resources, preventing their application to processes on geological time-scales. Eco-evolutionary models have been broadly adopted by the marine and terrestrial ecology communities, but have been largely ignored by earth-systems or biogeochemical modellers. Most global-scale ecosystem models are parameter rich, but represent only a miniscule fraction of biodiversity as large parts of the ecosystem are modelled using a single functional group parametrized from growth models of a single species. Such models are unable to replicate realistic ecosystem-level functional responses to environmental conditions (Franks, 2009), which show dramatic deviations from the typically hyperbolic functional responses of single species in the laboratory. Eco-evolutionary models have been able to predict the disparities between laboratory and ecosystem growth rates as functions of a variety of different environmental conditions, including the response of phytoplankton to nutrient levels (Lomas et al., 2014) and temperature levels (Sherman et al., 2016). Lomas (Lomas et al., 2014) used both adaptive dynamics and random sampling to study whole-community phosphorus uptake rates by phytoplankton in marine environments (Lomas et al., 2014). Using a model with cell radius as the primary trait, Lomas showed that the optimal cell size changed over a range of environmental conditions, leading to nutrient uptake kinetics that varied considerably as a function of environmental conditions. Emergent uptake rates diverged substantially from Michaelis-Menten kinetics, and fit observations of whole-community phosphorus uptake in the Sargasso Sea. The success of this approach suggests that it is possible to derive greatly simplified equations for the dynamics of the ecosystem and environment that still implicitly incorporate adaptive processes (Merico et al., 2009). Incorporating these ideas into global-scale earth-systems models of marine ecosystems would lead to more robust biological models that are not only simpler, but which better replicate the properties of real ecological communities.

Trait-based models, moment-closure techniques, and adaptive dynamics have made it possible to use complex adaptive systems theory to understand emergent patterns in ecosystems. As the adoption of these theories increases, the application of these methods is likely to become limited by our knowledge of physiology, and in particular the physiological trade-offs among different traits. Despite the recent rennaisance in trait-based modelling, nearly every important study using trait-based models of marine organisms uses parameters collected decades ago. New collaborations between experimentalists and modellers will be needed to fill in the large gaps in our knowledge. There are many challenges to understanding the full extent of phenotypic diversity in the biosphere, as the vast majority of organisms are still impossible to study in a laboratory setting. Despite these challenges, advances in biology, ranging from the invention of automated, high-throughput experimental techniques to advanced analytical methods for combining physiology and phylogeny (Bruggeman, 2011), point towards a bright future for the development of new, more accurate trait-based models.

## 3 Critical Transitions

The dynamics of complex adaptive systems are driven by non-linear interactions between small and large scale structures (Levin, 1998), giving rise to multiple attractors and the potential for system flips (May, 1977, Scheffer, 2009). Critical transitions between alternative stable states are an important part of the dynamics of real ecological systems (Holling, 1973), and they have become increasingly common as human activities have encroached on the environment, leading to potentially irreversible losses of key ecosystem services in both marine and terrestrial ecosystems. Recent advances in our understanding of critical transitions in ecological systems have been driven by two primary means: the adoption and invention of mathematical techniques to classify bifurcations and calculate the effect of interactions on multiple space and time scales and in complex network topologies, and the development of tools to model the coupled interactions between human behavior and ecological systems. The development of the field of social-ecological systems has led to a renaissance of interdisciplinary research which has spread ecological concepts such as resilience and robustness to social systems and brought ideas from game theory, economics, and optimal control theory to ecosystems management, enabling the solution of challenging problems at the interface between human and natural systems for the first time.

A critical transition is a sudden, large-scale change in the state of a system that occurs as the result of a small change in external conditions. Critical transitions occur at bifurcation points, where an equilibrium state or stable attractor vanishes or loses stability, causing the system to jump to an alternative stable state. Systems driven across critical transitions show *hysteresis*: reversal of the environmental stimulus does not return the system to the original state, as the basin of attraction for the alternative stable state persists even when the stability of the original state is restored. Alternative stable states are a general consequence of non-linearity, and can be expected when system components have threshold behaviors. For example, anoxia in the sub-surface waters of marine and lacustrine ecosystems can trigger the release of sedimentary phosphorus, leading to a self-sustaining process in which the released phosphorus enhances primary productivity, maintains oxygen depletion, and leads to a complete reorganization of the food web (Scheffer and van Nes, 2004, Van Cappellen and Ingall, 1994). Similar types of thresholds led to alternative stable states in tropical forest and grassland ecosystems (Staver et al., 2011a,b), where fire is the primary control on tree growth but is inhibited from spreading when tree cover exceeds a certain threshold, or in coral ecosystems, where grazer control of macro-algae can be lost at high macro-algal density, eventually leading to the domination of macro-algae over corals (Knowlton, 1992). Critical transitions are believed to be the cause of coastal dead zones (Diaz and Rosenberg, 2008) and the collapse of important fisheries (Sanchirico and Wilen, 2007), and there is cause to be concerned that anthropogenic climate change may trigger additional, high-impact critical transitions in the near future (Gattuso et al., 2015, Lenton et al., 2009).

Once a potential critical transition has been identified, it becomes important to quantify the *resilience* and *robustness* of the alternative ecosystem states. The terms resilience and robustness are defined in different ways in different disciplines (Levin and Lubchenco, 2008). Depending on the context they can refer to the size of the basin of attraction of a given state, or the time-scale for the decay of perturbations away from a stable state. Ecological resilience is crucial for the management of ecological systems, which should promote the resilience of desirable states and reduce the resilience of undesirable ones (Levin et al., 2013). Resilience and robustness are straightforward to calculate in simple models of critical transitions based on bifurcations, but the high-dimensionality of ecological systems is a serious obstacle to the application of criteria based on resilience and robustness to real systems. In order to overcome these obstacles, ecologists have used methods that allow for computation across multiple space and time scales, borrowing them from statistical physics and non-linear dynamics or developing them from scratch. As a result, we can now study the resilience and robustness of processes occurring in inherently high-dimensional systems or on complex networks.

Bifurcation theory has been used to develop early warning indicators for impending critical transitions from time-series data (Scheffer et al., 2012). Near a critical transition, system dynamics are governed by the normal form of the differential equation at the critical transition point. Disparate systems share the same normal forms, and thus exhibit universal behavior during critical transitions. The following class of models has been used to describe the linearized dynamics of a system that is approaching a saddle-node bifurcation (Strogatz, 2014), which is the most commonly observed critical transition in ecological systems:

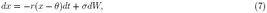

where *W* is the Wiener process (Gardiner et al., 1985) and *r* is assumed to slowly approach 0 as a function of the time *t*. The variance of this process is 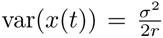, and the autocorrelation is 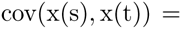 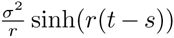, for *s* < *t*. As the bifurcation point is approached, the relaxation time towards steady state diverges, which is a process known as critical slowing down and which is reflected in the singular behavior of the statistical moments of the time series. These formulas suggest that increases in these statistical moments indicate the loss of resilience and an impending critical transition. Early warning indicators for critical transitions have been constructed using the autocorrelation (Held and Kleinen, 2004), variance (Carpenter and Brock, 2006), skewness (Guttal and Jayaprakash, 2008), or from statistical estimation of the parameters in equation 7 (Boettiger and Hastings, 2012b), all calculated from time-series data. Verifying the efficacy of early warning signals is a difficult statistical problem, as most systems of interest are too large and slowly evolving to admit experimental manipulations. As a result there is concern that commonly used early warning indicators generate high false-positive rates (Boettiger and Hastings, 2012a, Lenton, 2011). These challenges may be alleviated by adopting a model-based approach, treating each individual system or transition as a separate case, and using the methods presented in section 2 to derive reduced models from the small-scale interactions present in the system under study.

We have discussed critical transitions in the context of spatially homogeneous, well-mixed systems, but many real ecological systems are spatially heterogeneous and have mixing and dispersal time-scales that are longer than the time-scale of local ecological competition. To understand how spatial heterogeneity alters resilience requires methods that can bridge from the local scale dynamics to global-scale patterns. Numerical simulations (van Nes and Scheffer, 2005) and theoretical calculations using statistical physics (Martín et al., 2015) suggest that spatial heterogeneity smooths critical transitions, which appear as *first-order phase transitions* in spatially explicit systems. Heterogeneity promotes the preservation of local domains dominated by each possible equilibrium state, leading to a gradual, *second-order phase transition* as environmental parameters change. The interactions between diffusivity, heterogeneity, and alternative stable states can generate complex spatial patterns, including fractal structures and Turing patterns (Bak et al., 1987, Turing, 1952). Spatial heterogeneity has been observed to enhance resilience in numerous real ecological systems, including the tent caterpillar of British Columbia (Wellington, 1964), which is able to persist in isolated patches when mean environmental conditions would not favor its survival. These results suggest that artificially introducing heterogeneity, through reserves or environmental modification, or reducing diffusivity by inhibiting the movement or dispersal of ecosystem elements, might increase the resilience of certain ecosystem states.

The smoothing of first-order phase transitions by spatial heterogeneity is one of many mechanisms that generate *critical phenomena*, which are the scale-free, power law behaviors of spatial correlations functions and statistical moments at a second-order phase transition. Power laws are common in ecological systems, and can be observed in the spectral distribution of different types of organisms and habitats as well as in the magnitude and abundance of different types of events. Statistical physics classifies critical phenomena using their scaling exponents, with each set of scaling exponents defining a universality class, allowing for the derivation of mechanistic models using exponents measured from data. The concept of criticality has been expanded in scope to describe ecological phenomena. Unlike typical systems from physics, ecological systems can self-organize to their critical point, robustly exhibiting critical behavior without requiring the tuning of any parameters. Self-organized criticality has been used to describe two broad classes of ecological phenomena (Pascual and Guichard, 2005); including disease dynamics, which are described using percolation theory; and the dynamics of localized disturbances, which has been used to describe the interaction between fires and vegetation in a variety of ecosystems. The latter set of systems fall into a well-defined set of ecological universality classes, exhibiting power-law scaling that is insensitive to model parameters (Pascual et al., 2002). This observation has made it possible to diagnose the mechanisms driving pattern formation by examining data: phytoplankton patchiness is likely driven by ocean turbulence rather than ecological interactions, and similar patterns in vegetative ecosystems are likely to be driven by ecological interactions. Improvements in data collection are allowing for increased precision in the calculation of power laws, which will help resolve controversies about the prevalence of power laws in nature and their potential overuse in descriptions of natural phenomena (Clauset et al., 2009). Criticality is also appealing from an evolutionary perspective, and systems at critical poins maximize their sensitivity to environmental stimuli. Thus it has been proposed that criticality is an evolutionary attractor for both animal groups and neural systems (Mora and Bialek, 2011). Studies of criticality in ecological systems will be aided by the continued improvement in both theoretical and empirical methods.

The investigation of resilience has led ecologists to the study of network structures (Montoya et al., 2006). Networks arise naturally in complex adaptive systems, representing spatial structures, such as larval dispersal networks on coral reefs or island ecosystems, or the structure of local interactions between distinct elements, as in food webs or (human or animal) social networks. Network theoretic techniques were developed to understand and quantify complex systems, and although network theory is still a new field, it has had a profound effect on conservation science, epidemiology, and even economic regulation (Newman, 2003). The analytical techniques used to study the networks that arise in complex adaptive systems mirror those introduced in section 2, borrowing from equilibrium and non-equilibrium statistical physics to calculate the global implications of local network structures and interactions. One of the most important network features for resilience and robustness is modularity (or its opposite, connectivity) (May et al., 2008). Highly connected networks can have a homogeneous response to perturbations, suppressing rare patches of the alternative state through non-linear feedbacks. As a result, highly connected networks can be prone to critical transitions. Modular networks contain isolated sub-networks which can act as reservoirs of a state or as barriers against contagion, promoting a smooth response to external perturbations and adaptability at the global level. Under certain circumstances, such as heterogeneity in the dispersal rate of different system components, connectivity can enhance resilience by allowing the promoters of the desirable state to spread widely. Balancing these trade-offs has been an important consideration in the design of no-take areas and marine protected areas in coral ecosystems: denser spacings between reserves allow for more rapid replenishment of locally depleted populations, but also increase the global exposure to common risks (Almany et al., 2009). Food webs networks may be an exception to these rules, as studies have shown that increased connectance decreases the probability of large-scale species loss when links are randomly removed (Dunne et al., 2002), which is likely due to differences between the contagion-like processes that occur on networks that represent spatial structure and the competitive interactions occurring on food webs. Highly connected food webs may be better able to re-organize after the loss of a node, maintaining the flow of carbon and nutrients through the ecosystem. Connectance can thus enhance resilience by increasing the redundancy of function in a food web. The analysis of network structure in food webs has also helped identify *keystone species*, which are food web members with low numerical abundance but an outsized impact on food web dynamics, resilience, and robustness (Paine, 1969). Interdisciplinary research combining metagenomics, network theory, and ecology has transformed food web reconstruction and analysis from a labor-intensive process only applicable to a small number of ecosystems into a high-throughput process that can infer the ecological interactions between unculturable microbial species (Kurtz et al., 2015), allowing for unprecedented tests of ecological theories in microbial contexts. Algorithms have been invented to automatically identify keystone species (Berry and Widder, 2014), which in the case of the human microbiome has led to novel treatment regimes that are able to restore ecosystem resilience after antibiotic usage Buffie et al. (2015).

Human activity provides the environmental perturbation driving many of the critical transitions observed in nature, and feedbacks between human behavior and environmental conditions are therefore an important part of the dynamics of critical transitions. Social-ecological systems, which combine the dynamics of human and natural systems, are complex adaptive systems (Levin et al., 2013). In these systems, the adaptation of human behavior to local conditions can be modeled analogously to eco-evolutionary processes: the environment causes selection on human behavior, which feeds back to change the environment. Social-ecological systems must be studied in an interdisciplinary way, using models of human behavior that come from game theory or economics in combination with environmental models derived from ecology. The study of social-ecological systems has helped elucidate an important aspect of interactions between humans and their environment: there is often a time-scale separation between the ecological dynamics and the social dynamics. Profit maximizing humans are capable of rapid adaptation and can constantly optimize their behavior, but the environment may respond on a much slower time-scale, masking a dangerous loss of resilience at the ecosystem level. Multi-scale methods and singular perturbation theories have been applied to these problems to understand the interactions between the disparate social and ecological time-scales. These dynamics lead to *adaptive-cycles* (Gunderson, 2001, Holling, 1986), in which long exploitation and conservation phases are punctuated by rapid collapse and reorganization phases. An inability for management to recognize the slow-dynamics in social-ecological systems has been implicated as the cause of critical transitions in lakes, in which the fast variable is fertilizer usage and the slow variable is sediment phosphorus, and grazing-lands, in which the fast variable is grazing pressure and the slow variable is woody vegetation cover (Carpenter et al., 2001). Here the transition of human behavior from exploitative to protective occurs only after ecosystem collapse, causing the system to become trapped in an undesirable stable state for a long period of time. Using singular perturbation theory it is often possible to split the dynamics of social-ecological systems into fast and slow variables, reducing the system to the dynamics of the slow-variables on a low-dimensional slow-manifold (Fenichel, 1979). This ansatz has allowed for the rigorous analytical treatments of complex social-ecological models, allowing for the calculation of optimal management strategies for systems with alternative stable states such as coral reefs (Crépin, 2007). This work showed that so-called optimal fisheries management could trigger critical-transitions depending on the initial state of the coral-reef and the utility of the fish stocks, implying the importance of careful consideration of the societal discount rates in the determination of optimal management strategies.

Important challenges remain for the use of the complex adaptive systems to understand critical transitions, robustness, and resilience. Robustness and resilience are emergent properties of systems, arising from eco-evolutionary or social-ecological dynamics over long time scales. Understanding why either robust or fragile structures emerge in financial organizations, microbial systems, or marine food webs would be of direct utility to physicians, scientists, and regulators working to prevent catastrophic change. Though little is known about this subject, early efforts to understand the evolution of food web structures using numerical simulations (Drossel et al., 2004) and statistical physics (Bunin, 2016) are encouraging. Phylogenetic diversity is believed to play a role in the resilience of ecosystems to critical transitions (Norberg, 2004), but it has been difficult to make a quantitative link between diversity and adaptive capacity in real ecosystems at risk for critical transitions. Efforts are underway to understand the effects of commercial fishing on a wide variety of marine species (Pinsky and Palumbi, 2014), to determine whether observed physiological changes are due to plastic responses or rapid evolution. These studies suggest significant diversity loss in overfished populations, with uncertain consequences for marine fisheries (Eikeset et al., 2013). Incorporating diversity into models of critical transitions is a key step to unlocking the full potential of the complex adaptive systems approach. The study of emergent patterns has been successful in systems where it has been possible to quantify the trade-offs between different ecological strategies and to solve for ecological winners as a function of environmental conditions, but such simplification is unavailable when human behavior is governed by complex game-theoretic dynamics or strategic interactions between agents (Levin et al., 2013), and new techniques will be needed to study social-ecological interactions in these more complex cases. One of the primary goals of the study of critical transitions is to make actionable recommendations about the management of ecological systems, but the reliability of any recommendations is fundamentally limited by the large uncertainty surrounding ecological models. Improper treatment of model uncertainty is particularly risky when critical transitions are possible, as the cost of an inadvertent system flip may be exorbitant. Thus proper uncertainty quantification is an important step in development of management strategies for critical transitions (Polasky et al., 2011).

## 4 Public Goods

Public goods, common pool resources, and cooperation are universal features of complex adaptive systems, with each playing important roles in a wide variety of systems, from microbial ecosystems to human societies. Understanding the conditions that promote or inhibit cooperative behavior is thus an important problem, whose solution would help inspire the development of strategies for solving complex challenges, such as curing cancer, negotiating robust international agreements, or calculating the amount of organic carbon sequestered in the ocean. Recent developments have been catalyzed by the framing of public goods problems in the context of complex adaptive systems. Advances in evolutionary theory and game theory have elucidated the importance of spatial structure in contributing to the evolution of cooperation. Progress has been particularly promising in microbial ecosystems, which have been explored using new high-throughput techniques, revealing rich microscale dynamics involving complex metabolic interactions, public goods production, and competition between producers and scroungers. Improvements in high-performance computing have helped the modeling of systems that are driven by highly heterogeneous microscopic dynamics, making use of new techniques that couple multi-agent ecological processes with physical processes occurring on multiple scales. These insights into the evolution of cooperation in the natural world will help us not only to understand the dynamics of important microbial processes, but also to create stable mechanisms for promoting cooperation in human societies and the sustainable use of resources.

A fundamental problem in evolutionary theory that apparently has challenged theorists since Darwin is to understand how natural selection can favor individuals which sacrifice their fitness for the benefit of others. This can be seen by considering evolutionary dynamics within simple games such as the *prisoner’s dilemma or the public goods game* (Axelrod, 2006). In the latter, there are two phenotypes, a *producer* and a *scrounger*. The producer pays a cost *c* to give a benefit *b* to other individuals in the population, while a scrounger pays no cost but still receives the benefits from the producers. In a well-mixed population without any heterogeneity or structure in the interactions, the scroungers will dominate the producers and drive them to extinction, lowering the fitness of the entire community and leading to the *tragedy of the commons*. In order for public goods production to be sustainable, structure is required to allow producers to benefit more from production than scroungers. Evolutionary theorists have identified many such mechanisms, which usually rely on the spatial assortment of similar individuals (Damore and Gore, 2012). In structured environments such as a bacterial biofilm (Nadell et al., 2008), producers of the public good will be clustered near their kin and near other producers, allowing producers to receive the majority of the benefits from continued public goods production.

The simplest type of structure that can sustain cooperation is one in which the producer of a public good receives a greater benefit from production than neighboring cells. As an example, public goods are often molecules that are produced within a cell and released into the environment, reaching their highest concentration near their source of production. Leaky metabolic functions are a common source of public goods benefiting the producer more than the scroungers. Examples include marine nitrogen fixation and hydrogen peroxide detoxification: in the former case some percentage of fixed nitrogen escapes from the inside of the cell, and in the latter case hydrogen peroxide metabolism inside phytoplankton cells consumes the majority of hydrogen peroxide in the water column. The benefits of leaky processes are non-linear and hence frequency dependent: the relative benefit to scroungers increases at high producer population density. This dynamic is captured by the *snowdrift game*, in which players trapped by a hypothetical snowdrift must decide whether or not to expend e ort to help shovel the snow. If both players shovel, then each receives a score of *b* – *c*/2, reflecting their shared effort. If only one player shovels, then that player receives *b* − *c* and the other player *b*, and both players score 0 if neither decides to shovel. This game differs from the prisoner’s dilemma because there is no dominant strategy: the best move is to play oppositely from one’s opponent. The snowdrift game has been used to model the cooperation and competition between yeast cells that with different sucrose utilization capabilities (Gore et al., 2009). Some yeast cells produce an enzyme called *invertase*, which allows them to convert sucrose into fructose and glucose. The benefits of invertase production are leaky, and the majority of the sugars are lost from the cell due to diffusion, and can be taken up by cells that do not produce invertase. Because nutrient uptake is a nonlinear function of concentration, public goods production is advantageous when producers are rare and disadvantageous when they are common. This negative frequency dependent selection leads to the coexistence of producers and scroungers, and model experiments are very well fit by simulations of the snow-drift game.

The study of spatial structure has helped reveal the circumstances under which public goods are produced in the marine environment. Heterotrophic bacteria and phytoplankton rely on extracellular enzymes and ligands to enhance nutrient availability, either by directly binding insoluble nutrients such as iron or by liberating monomers from large organic molecules. Extracellular enzyme activity is both commonly detected in the ocean and critically important for numerous biogeochemical processes, but the factors controlling the release of enzymes are poorly understood. In particular, it is nearly impossible to discriminate between enzymes that are attached to the cell-surface and those that are freely difiusing, leading to uncertainty about the effect of these compounds on nutrient fields. Traving (Traving et al., 2015) and Vetter (Vetter et al., 1998) model the dynamics of extracellular enzymes in both open and confined marine environments via advection diffusion equations:

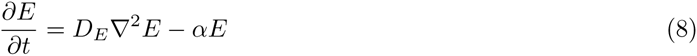

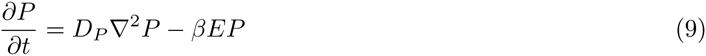

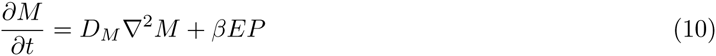

Here *E* is the enzyme concentration, *P* is the substrate concentration, *M* is the momoner concentration, *α* is the enzyme decay rate, *β* is the reaction rate constant for enzyme activity, and the coefficients *D* are diffusivities. Monomer uptake and enzyme production are represented by boundary conditions at the cell surface, and nutrient supply is represented by setting *P*, *M*, and *E* to constant values at infinity. The limit *D*_*E*_ = 0 corresponds to the strategy of attached enzymes. These equations can be solved at equilibrium, allowing for the calculation of extracellular nutrient fields and the rate of flux of monomers to the cell surface. These calculations showed that enzyme diffusivity has an enormous impact on the flux of nutrients to the cell and on the external nutrient field. Attached enzymes are the most cost-effective strategy for a single cell, producing the highest per-enzyme nutrient acquisition rates. As the diffusivity of enzymes increase, the realized nutrient flux drops but the total nutrient production increases, potentially benefiting cells near the source of enzyme production. These findings are consistent with observations suggesting that extracellular enzyme release is restricted to environments with a high degree of spatial structure, such as sinking particles or microbial biofilms.

Bacterial biofilms are complex structures whose properties are determined by microscopic biophysical interactions. These interactions can control the sustainability of the production of different types of public goods by altering the diffusivity of solutes and the relatedness of nearby organisms. The theory of complex adaptive systems has inspired the development of a powerful paradigm for modeling biofilms, integrating the interactions between the behavior of individual cells, fluid flow, and evolutionary dynamics (Nadell et al., 2013). Spatial structure is dictated by the fluid mechanical properties of the biofilm, and consideration of the characteristic scales of the biofilm, such as population density, thickness, external nutrient supply rate, or Reynolds number, can be used to de ne different biofilm states. For instance, the ratio of nutrient supply to nutrient demand describes the extent to which bacterial growth is confined to the outer layers of a biofilm (Nadell et al., 2010, Picioreanu et al., 1998), and the product of cellular surface area, the per-unit fitness enhancement produced by the public good, and the reciprocal of the enzyme diffusivity measures the extent to which public goods production benefits the individual versus the collective (Driscoll and Pepper, 2010), which can be used to derive an explicit criterion for the evolution of public goods production in microbial biofilms. These dimensionless quantities are emergent properties of the biofilm, and eco-evolutionary forces can cause them to change substantially. *Vibrio cholerae* communities, for example, can insulate themselves against invasion by scroungers through the production of extracellular polymers which anchor mother and daughter cells together (Nadell et al., 2015), resulting in highly structured biofilms that favor the production of public goods such as the *chitinase* enzyme. This modeling framework has been further enabled by the development of high-performance computing and in particular individual based models (Grimm and Railsback, 2013), which allow computer simulations to track every cell in a model biofilm. Such simulations have been used to show the emergence of spatial lineage segregation in nutrient limited biofilms, which favors the evolution of public goods by separating producers from scroungers. High resolution measurements have since confirmed the presence of a such a transition in real biofilms (Momeni et al., 2013), and simulation studies have become indispensable tools for identifying important mechanisms in biofilms.

Biofilms are ideal testbeds in which to investigate the evolution of cooperation and the production of public goods. Until recently it has been impossible to perform high-resolution observations of social interactions within microbial biofilms, but this has been changed by the development of a variety of new techniques such as high-throughput, culture-free genomics, bioinformatics, and computer vision. For instance, microscopy has been used to image entire biofilms at single-cell resolution, revealing both their physical architecture and the spatial distribution of different phenotypes (which are tagged with flourescent markers). Combined with experimental manipulations, these techniques have led to the determination of how different types of mucus influence the material properties of the biofilm (Drescher et al., 2016). On a broader scale, genomic techniques have been used to study the distribution of producer and scrounger traits within field populations. Heterotrophic marine bacteria excrete siderophores and ligands, which public goods that bind iron, allowing it to be absorbed by cells and preventing it from precipitating out of solution. The distribution of genes for siderophore production and utilization within Vibronaceae colonies on sinking particles showed that siderophore producers faced competition from scroungers only on larger particles (Cordero et al., 2012). Non-producers were able to colonize particles which already had an established producer community, leading to a regular successional cycle in which bacterial community metabolism shifts from the use of particulate organic matter to the recycling of dissolved organic matter generated by the producer community (Datta et al., 2016). This may lead to competition for oxygen and nutrients and a decrease in the efficiency or organic matter remineralization. Bioinformatic studies of syntrophic microbial communities from the human gut microbiome showed that genes for production of different extracellular enzymes are strongly correlated with each-other, suggesting that public goods are produced by a specialized subset of community members (Rakoff-Nahoum et al., 2014). This pattern has been observed in a wide-variety of distinct microbial ecosystems, which has lead to the proposal that gene loss is more likely to benefit non-producers than producers, leading different linearges to either specialize in public goods production or cheating (Morris et al., 2012). This generic mechanism is known as the *black queen hypothesis*, and it has been used to explain the evolution of metabolic dependencies throughout microbial ecosystems.

Public goods and cooperation are important features of both human societies and microbial ecosystems, and there is optimism that insights derived from cooperation in biological systems can be used to devise mechanisms that promote cooperation in human societies (Levin, 2014). Although there are fundamental differences between cooperation in human systems and natural systems, evolutionary principles determine the stability of cooperation in both types of systems (West et al., 2011). Despite the shared Darwinian framework, the transfer of ideas between social and biological evolution has been slow, and top-down efforts to regulate the use of public goods and common pool resources fail as often as they succeed. Management strategies involve making predictions about how human beings will respond to rules and regulations, and uncertainty in these responses has complicated efforts to successfully manage resource (Fulton et al., 2011). For instance, many proposed solutions to common pool resource problems in human societies have relied on an overly simplistic rational-actor model, which suggests that common pool resources need to either be monopolized by the government or privatized entirely (Dietz, 2005). Although there are numerous examples examples of long-lasting social institutions that have sustainably managed common pool resources, extreme approaches like *laissez-faire* principles or central planning are not found amongst them. These successes have taught us a number of important lessons about managing public goods, leading to the identification of several important design principles shared by the majority of long-lived cooperative societies (Ostrom, 2015). Darwinian thinking favors a polycentric approach over a global approach, promoting cooperation by focusing on small-scales, and encouraging experimentation with a diverse series of local strategies so that heterogeneities between regions can be overcome (Ostrom, 2010). The theory of complex adaptive systems provides tools that can be used to understand and design institutions that promote cooperation by enabling the consideration of multiple scales of interaction as well as socio-evolutionary dynamics. The acceleration of human activity since the industrial revolution threatens common pool resources essential to our well-being and survival, and the resulting challenges will be amongst the most important and exciting problems faced by social scientists and governments in the next century.

## 5 Discussion

Within the past twenty years, the science of complex adaptive systems have evolved from an abstract concept into a set of powerful tools for solving real problems. Central to this transformation has been the development of methods for studying the interactions among structures at different scales, which have helped reveal the causes of ecological patterns. Progress has been driven by interdisciplinary collaborations: moment closures, phase transitions, and high performance computing have been adapted to ecological problems and are indispensable for understanding critical phenomena and the emergence of macroscale structure from microscale interactions. Nonlinear dynamics and stochasticity have been emphasized, and ideas from probability theory and dynamical systems have contributed to the identification of tipping points and quantification of the resilience of ecological systems to external perturbations. The complex adaptive systems framework has also inspired the invention of new techniques. Adaptive dynamics and trait-based models represent a first step towards a statistical mechanics of types (Levin, 2003), allowing for the coupling of ecological and evolutionary dynamics by accounting for the appearance of novel types. Network theory has helped introduce realism into descriptions of interactions between individuals or ecosystem components, which at first were too closely inspired by physical systems, leading to the development of new tools for calculating dynamics across scales in systems with complex interactions. These theoretical advances have arose alongside a revolution in data collection and molecular ecology, allowing for the inference of the complex network of interactions within ecosystems, and revealing mechanisms which maintain cooperation and ensure the production of critical public goods.

There are several important challenges that must be overcome to realize the full potential of complex adaptive systems: we need a better understanding of the inherent trade-offs between different ecological strategies, new methods must be developed to study emergence in systems with complex species interactions, and we must build tools to help regulate the use of public goods and common pool resources. Complex adaptive systems are characterized by the appearance of novel types, which must be parametrized to formulate a closed system. Trait-based models accomplish this task by utilizing the physiological trade-offs between different traits, and our lack of understanding of key physiological trade offs is a fundamental limit on further progress. Despite this, many state-of-the-art trait-based models make use of physiological measurements that are over thirty years old. Laboratory techniques have improved considerably since then, and renewed collaborations between physiologists and theorists could lead to dramatic improvements of eco-evolutionary models. Eco-evolutionary models have been tremendously successful in shaping our understanding of simple ecosystems, such as those whose dynamics are dominated by trees (Enquist et al., 2015), phytoplankton (Litchman and Klausmeier, 2008), or other primary producers. When only a small number of trophic levels are present or when the food web has a simple structure, the evolutionary dynamics can be simplified: in marine ecosystems phytoplankton quickly reach the optimal strategy for a given environment, collapsing the evolutionary dynamics onto a slow manifold and causing the trait-distribution to track the slow evolution of the environment. Such simplifications are unavailable in more complex communities, such as microbial biofilms attached to marine snow or within the human microbiome, as the complex interactions of these communities prevent the simple calculation of optimal strategies. Some promising new techniques have appeared in recent years, such as community flux balance analysis (Khandelwal et al., 2013), maximum entropy production (Vallino, 2010), or cavity methods from statistical physics (Bunin, 2016), but much further works is required to apply these tools to understand real systems.

Perhaps the most important challenge facing ecosystems science is understanding how to manage the interactions of humans with their environment (Dietz et al., 2003). The central features of complex adaptive systems, such as alternative stable states or conflicts between individual goals and the social optimum, present common pitfalls to the management of natural resources and social-ecological systems. Despite the importance of these features, management is still typically guided by models that exclude non-linearities or the possibility for evolution and adaptation. As a result, the plans of management are often foiled by human behavior and the complexity of the underlying ecosystem dynamics, leading to critical transitions and the irreversible collapse of key ecosystem services. Although many basic mechanisms that lead to the failure of traditional management strategies are known, more work needs to be done before the complex adaptive systems framework can be fully adopted as a tool for managing ecosystems. As the Earth becomes more crowded and ecosystems become more stressed, the fate of future generations will rely on how well we manage and secure the public goods and common pool resources contained in our ecosystems.

## 6 Acknowledgments

Simon Levin acknowledges funding from the NSF Dimensions of Biodiversity grant OCE-1046001, NSF grant GEO-1211972, NSF grant OCE-1426746, ARO grant W911NF-11-1-0385, ARO grant W911NF-14-1-0431 and by the Nordforsk-funded project Green Growth Based on Marine Resource: Ecological and Socio-Economic Constraints (GreenMAR). George Hagstrom acknowledges funding from NSF Dimensions of Biodiversity grant OCE-1046001, ARO grant W911NF-14-1-0431, and ARO grant W911NF-11-1-0385. The authors declare that they have no conflicts of interest.

## References

Almany G, Connolly S, Heath D, Hogan J, Jones G, McCook L, Mills M, Pressey R, and Williamson D. 2009. Connectivity, biodiversity conservation and the design of marine reserve networks for coral reefs. Coral Reefs 28(2): 339–351.

Andersen KH, Berge T, Gonçalves R, Hartvig M, Heuschele J, Hylander S, Jacobsen NS, Lindemann C, Martens EA, Neuheimer AB, et al. 2016. Characteristic sizes of life in the oceans, from bacteria to whales. Marine Science 8.

Anderson PW. 1972. More is different. Science 177(4047): 393–396.

Armstrong RA. 1994. Grazing limitation and nutrient limitation in marine ecosystems: steady state solutions of an ecosystem model with multiple food chains. Limnology and Oceanography 39(3): 597–608.

Arrow KJ, Cropper ML, Gollier C, Groom B, Heal GM, Newell RG, Nordhaus WD, Pindyck RS, Pizer WA, Portney PR, et al. 2014. Should governments use a declining discount rate in project analysis? Review of Environmental Economics and Policy p. reu008.

Arthur WB. 1994. Increasing returns and path dependence in the economy. University of Michigan Press.

Axelrod RM. 2006. The evolution of cooperation. Basic books.

Bak P, Tang C, and Wiesenfeld K. 1987. Self-organized criticality: An explanation of the 1/f noise. Physical review letters 59(4): 381.

Berdahl A, Torney CJ, Ioannou CC, Faria JJ, and Couzin ID. 2013. Emergent sensing of complex environments by mobile animal groups. Science 339(6119): 574–576.

Berry D and Widder S. 2014. Deciphering microbial interactions and detecting keystone species with co-occurrence networks. Frontiers in microbiology 5: 219.

Bialek W, Cavagna A, Giardina I, Mora T, Silvestri E, Viale M, and Walczak AM. 2012. Statistical mechanics for natural flocks of birds. Proceedings of the National Academy of Sciences 109(13): 4786–4791.

Boettiger C and Hastings A. 2012a. Early warning signals and the prosecutor’s fallacy. Proceedings of the Royal Society of London B: Biological Sciences p. 2085.

Boettiger C and Hastings A. 2012b. Quantifying limits to detection of early warning for critical transitions. Journal of the Royal Society Interface 9(75): 2527–2539.

Bruggeman J. 2011. A phylogenetic approach to the estimation of phytoplankton traits. Journal of Phycology 47(1): 52–65.

Brush ER, Leonard NE, and Levin SA. 2016. The content and availability of information affects the evolution of social-information gathering strategies. Theoretical Ecology pp. 1–22.

Buffie CG, Bucci V, Stein RR, McKenney PT, Ling L, Gobourne A, No D, Liu H, Kinnebrew M, Viale A, et al. 2015. Precision microbiome reconstitution restores bile acid mediated resistance to clostridium difficile. Nature 517(7533): 205–208.

Bunin G. 2016. Interaction patterns and diversity in assembled ecological communities. arXiv preprint arXiv:1607.04734.

Carpenter SR and Brock WA. 2006. Rising variance: a leading indicator of ecological transition. Ecology letters 9(3): 311–318.

Carpenter SR, Ludwig D, and Brock WA. 1999. Management of eutrophication for lakes subject to potentially irreversible change. Ecological applications 9(3): 751–771.

Carpenter SR, Press MC, Huntly NJ, and Levin SA. 2001. Alternate states of ecosystems: evidence and some implications. In Ecology: achievement and challenge: the 41st Symposium of the British Ecological Society sponsored by the Ecological Society of America held at Orlando, Florida, USA, 10-13 April 2000., pp. 357–383.

Cavagna A, Cimarelli A, Giardina I, Parisi G, Santagati R, Stefanini F, and Viale M. 2010. Scale-free correlations in starling flocks. Proceedings of the National Academy of Sciences 107(26): 11865–11870.

Chapman S and Cowling TG. 1970. The mathematical theory of non-uniform gases: an account of the kinetic theory of viscosity, thermal conduction and diffusion in gases. Cambridge university press.

Clark JR, Lenton TM, Williams HT, and Daines SJ. 2013. Environmental selection and resource allocation determine spatial patterns in picophytoplankton cell size. Limnology and Oceanography 58(3): 1008–1022.

Clauset A, Shalizi CR, and Newman ME. 2009. Power-law distributions in empirical data. SIAM review 51(4): 661–703.

Cordero OX, Ventouras LA, DeLong EF, and Polz MF. 2012. Public good dynamics drive evolution of iron acquisition strategies in natural bacterioplankton populations. Proceedings of the National Academy of Sciences 109(49): 20059–20064.

Couzin ID and Krause J. 2003. Self-organization and collective behavior in vertebrates. Advances in the Study of Behavior 32: 1–75.

Couzin ID, Krause J, Franks NR, and Levin SA. 2005. Effective leadership and decision-making in animal groups on the move. Nature 433(7025): 513–516.

Crépin AS. 2007. Using fast and slow processes to manage resources with thresholds. Environmental and Resource Economics 36(2): 191–213.

Czirók A and Vicsek T. 2001. Collective motion. Fluctuations and scaling in biology pp. 177–242.

Daines SJ, Clark JR, and Lenton TM. 2014. Multiple environmental controls on phytoplankton growth strategies determine adaptive responses of the n: P ratio. Ecology letters 17(4): 414–425.

Damore JA and Gore J. 2012. Understanding microbial cooperation. Journal of theoretical biology 299: 31–41.

Darwin C. 1858. The Origin of Species by means of Natural Selection.

Datta MS, Sliwerska E, Gore J, Polz MF, and Cordero OX. 2016. Microbial interactions lead to rapid micro-scale successions on model marine particles. Nature Communications 7.

Diaz RJ and Rosenberg R. 2008. Spreading dead zones and consequences for marine ecosystems. Science 321(5891): 926–929.

Diekmann O et al. 2004. A beginner’s guide to adaptive dynamics. Banach Center Publications 63: 47–86.

Dietz T. 2005. The darwinian trope in the drama of the commons: variations on some themes by the ostroms. Journal of Economic Behavior & Organization 57(2): 205–225.

Dietz T, Ostrom E, and Stern PC. 2003. The struggle to govern the commons. science 302(5652): 1907–1912.

Drescher K, Dunkel J, Nadell CD, van Teeffelen S, Grnja I, Wingreen NS, Stone HA, and Bassler BL. 2016. Architectural transitions in vibrio cholerae biofilms at single-cell resolution. Proceedings of the National Academy of Sciences 113(14): E2066–E2072.

Driscoll WW and Pepper JW. 2010. Theory for the evolution of difiusible external goods. Evolution 64(9): 2682–2687.

Drossel B, McKane AJ, and Quince C. 2004. The impact of nonlinear functional responses on the long-term evolution of food web structure. Journal of Theoretical Biology 229(4): 539–548.

Dunne JA, Williams RJ, and Martinez ND. 2002. Network structure and biodiversity loss in food webs: robustness increases with connectance. Ecology letters 5(4): 558–567.

Eikeset AM, Richter A, Dunlop ES, Dieckmann U, and Stenseth NC. 2013. Economic repercussions of fisheries-induced evolution. Proceedings of the National Academy of Sciences 110(30): 12259–12264.

Enquist BJ, Norberg J, Bonser SP, Violle C, Webb CT, Henderson A, Sloat LL, and Savage VM. 2015. Chapter nine-scaling from traits to ecosystems: Developing a general trait driver theory via integrating trait-based and metabolic scaling theories. Advances in Ecological Research 52: 249–318.

Falkowski PG. 1997. Evolution of the nitrogen cycle and its influence on the biological sequestration of co 2 in the ocean. Nature 387(6630): 272–275.

Falkowski PG, Fenchel T, and Delong EF. 2008. The microbial engines that drive earth’s biogeochemical cycles. Science 320(5879): 1034–1039.

Fenichel N. 1979. Geometric singular perturbation theory for ordinary differential equations. Journal of Differential Equations 31(1): 53–98.

Filotas E, Parrott L, Burton PJ, Chazdon RL, Coates DK, Coll L, Haeussler S, Martin K, Nocentini S, and Puettmann KJ. 2014. Viewing forests through the lens of complex systems science. Ecosphere 5(1): 1–23.

Flierl G, Grünbaum D, Levin SA, and Olson D. 1999. From individuals to aggregations: the interplay between behavior and physics. Journal of Theoretical biology 196(4): 397–454.

Follows MJ, Dutkiewicz S, Grant S, and Chisholm SW. 2007. Emergent biogeography of microbial communities in a model ocean. science 315(5820): 1843–1846.

Franks PJS. 2009. Planktonic ecosystem models: perplexing parameterizations and a failure to fail. Journal of Plankton Research p. fbp069.

Fulton EA, Smith AD, Smith DC, and van Putten IE. 2011. Human behaviour: the key source of uncertainty in fisheries management. Fish and Fisheries 12(1): 2–17.

Galbraith ED and Martiny AC. 2015. A simple nutrient-dependence mechanism for predicting the stoichiometry of marine ecosystems. Proceedings of the National Academy of Sciences 112(27): 8199–8204.

Gardiner CW et al. 1985. Handbook of stochastic methods, volume 3. Springer Berlin.

Gattuso JP, Magnan A, Billé R, Cheung WWL, Howes EL, Joos F, Allemand D, Bopp L, Cooley SR, Eakin CM, Hoegh-Guldberg O, Kelly RP, Pörtner HO, Rogers AD, Baxter JM, Laffoley D, Osborn D, Rankovic A, Rochette J, Sumaila UR, Treyer S, and Turley C. 2015. Contrasting futures for ocean and society from different anthropogenic co2 emissions scenarios. Science 349(6243). URL http://science.sciencemag.org/content/349/6243/aac4722. http://science.sciencemag.org/content/349/6243/aac4722.full.pdf.

Geritz SA, Metz JA, Kisdi É, and Meszéna G. 1997. Dynamics of adaptation and evolutionary branching. Physical Review Letters 78(10): 2024.

Gore J, Youk H, and Van Oudenaarden A. 2009. Snowdrift game dynamics and facultative cheating in yeast. Nature 459(7244): 253–256.

Grimm V and Railsback SF. 2013. Individual-based modeling and ecology. Princeton university press.

Grünbaum D. 1994. Translating stochastic density-dependent individual behavior with sensory constraints to an eulerian model of animal swarming. Journal of mathematical biology 33(2): 139–161.

Grünbaum D. 1998. Schooling as a strategy for taxis in a noisy environment. Evolutionary Ecology 12(5): 503–522.

Grünbaum D and Okubo A. 1994. Modelling social animal aggregations. In Frontiers in mathematical biology, pp. 296–325. Springer.

Gunderson LH. 2001. Panarchy: understanding transformations in human and natural systems. Island press.

Guttal V and Jayaprakash C. 2008. Changing skewness: an early warning signal of regime shifts in ecosystems. Ecology letters 11(5): 450–460.

Hagstrom GI, Levin SA, and Martiny AC. 2016. Resource ratios and primary productivity in the ocean. In review http://biorxiv.org/content/early/2016/07/18/064543.

Hartvigsen G, Kinzig A, and Peterson G. 1998. Complex adaptive systems: Use and analysis of complex adaptive systems in ecosystem science: Overview of special section. Ecosystems 1(5): 427–430.

Hein A, Rosenthal SB, Hagstrom G, Berdahl A, Torney C, and Couzin I. 2015. The evolution of distributed sensing and collective computation in animal populations. eLife p. 10955.

Heino M. 1998. Management of evolving fish stocks. Canadian Journal of Fisheries and Aquatic Sciences 55(8): 1971–1982.

Held H and Kleinen T. 2004. Detection of climate system bifurcations by degenerate fingerprinting. Geo-physical Research Letters 31(23).

Holling CS. 1973. Resilience and stability of ecological systems. Annual review of ecology and systematics pp. 1–23.

Holling CS. 1986. The resilience of terrestrial ecosystems: local surprise and global change. Sustainable development of the biosphere pp. 292–317.

Jacob F. 1977. Evolution and tinkering. Science 196(4295): 1161–1166.

Katz Y, Tunstrøm K, Ioannou CC, Huepe C, and Couzin ID. 2011. Inferring the structure and dynamics of interactions in schooling fish. Proceedings of the National Academy of Sciences 108(46): 18720–18725.

Khandelwal RA, Olivier BG, Röling WF, Teusink B, and Bruggeman FJ. 2013. Community flux balance analysis for microbial consortia at balanced growth. PLoS One 8(5): e64567.

Kiørboe T. 1993. Turbulence, phytoplankton cell size, and the structure of pelagic food webs. Advances in marine biology 29: 1–72.

Klausmeier CA, Litchman E, Daufresne T, and Levin SA. 2004. Optimal nitrogen-to-phosphorus stoichiometry of phytoplankton. Nature 429(6988): 171–174.

Knowlton N. 1992. Thresholds and multiple stable states in coral reef community dynamics. American Zoologist 32(6): 674–682.

Kurtz ZD, Müller CL, Miraldi ER, Littman DR, Blaser MJ, and Bonneau RA. 2015. Sparse and compositionally robust inference of microbial ecological networks. PLoS Comput Biol 11(5): e1004226.

Lenton TM. 2011. Early warning of climate tipping points. Nature Climate Change 1(4): 201–209.

Lenton TM, Footitt A, Dlugolecki A, and Allianz Gruppe. 2009. Major tipping points in the earth’s climate system and consequences for the insurance sector. World Wildlife Fund.

Levin S. 2003. Complex adaptive systems: exploring the known, the unknown and the unknowable. Bulletin of the American Mathematical Society 40(1): 3–19.

Levin S, Xepapadeas T, Crépin AS, Norberg J, De Zeeuw A, Folke C, Hughes T, Arrow K, Barrett S, Daily G, et al. 2013. Social-ecological systems as complex adaptive systems: modeling and policy implications. Environment and Development Economics 18(02): 111–132.

Levin SA. 1992. The problem of pattern and scale in ecology: the robert h. macarthur award lecture. Ecology 73(6): 1943–1967.

Levin SA. 1998. Ecosystems and the biosphere as complex adaptive systems. Ecosystems 1(5): 431–436.

Levin SA. 2014. Public goods in relation to competition, cooperation, and spite. Proceedings of the National Academy of Sciences 111(Supplement 3): 10838–10845.

Levin SA, Barrett S, Aniyar S, Baumol W, Bliss C, Bolin B, Dasgupta P, Ehrlich P, Folke C, Gren IM, et al. 1998. Resilience in natural and socioeconomic systems. Environment and development economics 3(02): 221–262.

Levin SA and Lubchenco J. 2008. Resilience, robustness, and marine ecosystem-based management. Bio-science 58(1): 27–32.

Litchman E and Klausmeier CA. 2008. Trait-based community ecology of phytoplankton. Annual Review of Ecology, Evolution, and Systematics pp. 615–639.

Lomas MW, Bonachela JA, Levin SA, and Martiny AC. 2014. Impact of ocean phytoplankton diversity on phosphate uptake. Proceedings of the National Academy of Sciences 111(49): 17540–17545.

Margalef R, Miyares ME, and de Rubinat DBF. 1979. Functional morphology of organisms involved in red tides, as adapted to decaying turbulence. Elsevier/North-Holland.

Martín PV, Bonachela JA, Levin SA, and Muñoz MA. 2015. Eluding catastrophic shifts. Proceedings of the National Academy of Sciences 112(15): E1828–E1836.

Martiny AC, Pham CTA, Primeau FW, Vrugt JA, Moore JK, Levin SA, and Lomas MW. 2013. Strong latitudinal patterns in the elemental ratios of marine plankton and organic matter. Nature Geoscience 6(4): 279–283.

May RM. 1977. Thresholds and breakpoints in ecosystems with a multiplicity of stable states. Nature 269(5628): 471–477.

May RM, Levin SA, and Sugihara G. 2008. Complex systems: Ecology for bankers. Nature 451(7181): 893–895.

Merico A, Bruggeman J, and Wirtz K. 2009. A trait-based approach for downscaling complexity in plankton ecosystem models. Ecological Modelling 220(21): 3001–3010.

Messier C, Puettmann K, Chazdon R, Andersson KP, Angers VA, Brotons L, Filotas E, Tittler R, Parrott L, and Levin SA. 2015. From management to stewardship: viewing forests as complex adaptive systems in an uncertain world. Conservation Letters 8(5): 368–377.

Momeni B, Waite AJ, and Shou W. 2013. Spatial self-organization favors heterotypic cooperation over cheating. Elife 2: e00960.

Montoya JM, Pimm SL, and Solé RV. 2006. Ecological networks and their fragility. Nature 442(7100): 259–264.

Mora T and Bialek W. 2011. Are biological systems poised at criticality? Journal of Statistical Physics 144(2): 268–302.

Morris JJ, Lenski RE, and Zinser ER. 2012. The black queen hypothesis: evolution of dependencies through adaptive gene loss. mBio 3(2): e00036–12.

Mueller T and Fagan WF. 2008. Search and navigation in dynamic environments–from individual behaviors to population distributions. Oikos 117(5): 654–664.

Murray JD. 2002. Mathematical biology i: an introduction, vol. 17 of interdisciplinary applied mathematics.

Nadell CD, Bucci V, Drescher K, Levin SA, Bassler BL, and Xavier JB. 2013. Cutting through the complexity of cell collectives. In Proc. R. Soc. B, volume 280, p. 20122770. The Royal Society.

Nadell CD, Drescher K, Wingreen NS, and Bassler BL. 2015. Extracellular matrix structure governs invasion resistance in bacterial biofilms. The ISME journal 9(8): 1700–1709.

Nadell CD, Foster KR, and Xavier JB. 2010. Emergence of spatial structure in cell groups and the evolution of cooperation. PLoS Comput Biol 6(3): e1000716.

Nadell CD, Xavier JB, Levin SA, and Foster KR. 2008. The evolution of quorum sensing in bacterial biofilms. PLoS Biol 6(1): e14.

Newman ME. 2003. The structure and function of complex networks. SIAM review 45(2): 167–256.

Norberg J. 2004. Biodiversity and ecosystem functioning: a complex adaptive systems approach. Limnology and Oceanography 49(4part2): 1269–1277.

Norberg J, Swaney DP, Dushoff J, Lin J, Casagrandi R, and Levin SA. 2001. Phenotypic diversity and ecosystem functioning in changing environments: a theoretical framework. Proceedings of the National Academy of Sciences 98(20): 11376–11381.

Ostrom E. 2010. Polycentric systems for coping with collective action and global environmental change. Global Environmental Change 20(4): 550–557.

Ostrom E. 2015. Governing the commons. Cambridge university press.

Pacala SW and Silander J. 1985. Neighborhood models of plant population dynamics. i. single-species models of annuals. American Naturalist pp. 385–411.

Paine RT. 1969. A note on trophic complexity and community stability. The American Naturalist 103(929): 91–93.

Pascual M and Guichard F. 2005. Criticality and disturbance in spatial ecological systems. Trends in ecology & evolution 20(2): 88–95.

Pascual M, Roy M, Guichard F, and Flierl G. 2002. Cluster size distributions: signatures of self–organization in spatial ecologies. Philosophical Transactions of the Royal Society of London B: Biological Sciences 357(1421): 657–666.

Picioreanu C, Van Loosdrecht MC, Heijnen JJ, et al. 1998. Mathematical modeling of biofilm structure with a hybrid differential-discrete cellular automaton approach. Biotechnology and bioengineering 58(1): 101–116.

Pinsky ML and Palumbi SR. 2014. Meta-analysis reveals lower genetic diversity in overfished populations. Molecular Ecology 23(1): 29–39.

Polasky S, Carpenter SR, Folke C, and Keeler B. 2011. Decision-making under great uncertainty: environmental management in an era of global change. Trends in ecology & evolution 26(8): 398–404.

Rakoff-Nahoum S, Coyne MJ, and Comstock LE. 2014. An ecological network of polysaccharide utilization among human intestinal symbionts. Current Biology 24(1): 40–49.

Redfield AC. 1958. The biological control of chemical factors in the environment. American scientist 46(3): 230A–221.

Rodriguez-Iturbe I and Rinaldo A. 1997. Fractal river networks: chance and self-organization.

Rosenthal SB, Twomey CR, Hartnett AT, Wu HS, and Couzin ID. 2015. Revealing the hidden networks of interaction in mobile animal groups allows prediction of complex behavioral contagion. Proceedings of the National Academy of Sciences 112(15): 4690–4695.

Sanchirico JN and Wilen JE. 2007. Global marine fisheries resources: status and prospects. International Journal of Global Environmental Issues 7(2-3): 106–118.

Scheffer M. 2009. Critical transitions in nature and society. Princeton University Press.

Scheffer M, Carpenter SR, Lenton TM, Bascompte J, Brock W, Dakos V, Van De Koppel J, Van De Leemput IA, Levin SA, and Van Nes EH. 2012. Anticipating critical transitions. science 338(6105): 344–348.

Scheffer M and van Nes EH. 2004. Mechanisms for marine regime shifts: can we use lakes as microcosms for oceans? Progress in Oceanography 60(2): 303–319.

Sheffer E, Batterman SA, Levin SA, and Hedin LO. 2015. Biome-scale nitrogen fixation strategies selected by climatic constraints on nitrogen cycle. Nature plants 1: 15182.

Sherman E, Moore JK, Primeau F, and Tanouye D. 2016. Temperature influence on phytoplankton community growth rates. Global Biogeochemical Cycles 30(4): 550–559.

Shuter B. 1979. A model of physiological adaptation in unicellular algae. Journal of theoretical biology 78(4): 519–552.

Staver CA, Archibald S, and Levin SA. 2011a. The global extent and determinants of savanna and forest as alternative biome states. Science 334(6053): 230–232.

Staver CA, Archibald S, and Levin SA. 2011b. Tree cover in sub-saharan africa: rainfall and fire constrain forest and savanna as alternative stable states. Ecology 92(5): 1063–1072.

Steele JH. 1998. Regime shifts in marine ecosystems. Ecological Applications 8(sp1): S33–S36.

Strogatz SH. 2014. Nonlinear dynamics and chaos: with applications to physics, biology, chemistry, and engineering. Westview press.

Teng YC, Primeau FW, Moore JK, Lomas MW, and Martiny AC. 2014. Global-scale variations of the ratios of carbon to phosphorus in exported marine organic matter. Nature Geoscience 7(12): 895–898.

Toner J and Tu Y. 1995. Long-range order in a two-dimensional dynamical xy model: how birds fly together. Physical Review Letters 75(23): 4326.

Toseland ADSJ, Daines SJ, Clark JR, Kirkham A, Strauss J, Uhlig C, Lenton TM, Valentin K, Pearson GA, and Moulton V. 2013. The impact of temperature on marine phytoplankton resource allocation and metabolism. Nature Climate Change 3(11): 979–984.

Traving SJ, Thygesen UH, Riemann L, and Stedmon CA. 2015. A model of extracellular enzymes in free-living microbes: which strategy pays off? Applied and Environmental Microbiology 81(21): 7385–7393.

Tunstrøm K, Katz Y, Ioannou CC, Huepe C, Lutz MJ, and Couzin ID. 2013. Collective states, multistability and transitional behavior in schooling fish. PLoS Comput Biol 9(2): e1002915.

Turing AM. 1952. The chemical basis of morphogenesis. Philosophical Transactions of the Royal Society of London B: Biological Sciences 237(641): 37–72.

Tyrrell T. 1999. The relative influences of nitrogen and phosphorus on oceanic primary production. Nature 400(6744): 525–531.

Vallino JJ. 2010. Ecosystem biogeochemistry considered as a distributed metabolic network ordered by maximum entropy production. Philosophical Transactions of the Royal Society of London B: Biological Sciences 365(1545): 1417–1427.

Van Cappellen P and Ingall ED. 1994. Benthic phosphorus regeneration, net primary production, and ocean anoxia: A model of the coupled marine biogeochemical cycles of carbon and phosphorus. Paleoceanography 9(5): 677–692.

van Nes EH and Scheffer M. 2005. Implications of spatial heterogeneity for catastrophic regime shifts in ecosystems. Ecology 86(7): 1797–1807.

Vetter Y, Deming J, Jumars P, and Krieger-Brockett B. 1998. A predictive model of bacterial foraging by means of freely released extracellular enzymes. Microbial ecology 36(1): 75–92.

Vicsek T, Czirók A, Ben-Jacob E, Cohen I, and Shochet O. 1995. Novel type of phase transition in a system of self-driven particles. Physical review letters 75(6): 1226.

Viswanathan GM, Da Luz MG, Raposo EP, and Stanley HE. 2011. The physics of foraging: an introduction to random searches and biological encounters. Cambridge University Press.

Ward BA, Dutkiewicz S, Jahn O, and Follows MJ. 2012. A size-structured food-web model for the global ocean. Limnology and Oceanography 57(6): 1877–1891.

Ward BA and Follows MJ. 2016. Marine mixotrophy increases trophic transfer efficiency, mean organism size, and vertical carbon flux. Proceedings of the National Academy of Sciences 113(11): 2958–2963.

Wellington W. 1964. Qualitative changes in populations in unstable environments. The Canadian Entomologist 96(1-2): 436–451.

West SA, El Mouden C, and Gardner A. 2011. Sixteen common misconceptions about the evolution of cooperation in humans. Evolution and Human Behavior 32(4): 231–262.

Young GF, Scardovi L, Cavagna A, Giardina I, and Leonard NE. 2013. Starling flock networks manage uncertainty in consensus at low cost. PLoS Comput Biol 9(1): e1002894.

